# Heat-shock inducible CRISPR/Cas9 system generates heritable mutations in rice

**DOI:** 10.1101/418517

**Authors:** Soumen Nandy, Bhuvan Pathak, Shan Zhao, Vibha Srivastava

**Author notes:** Author for correspondence: 115 Plant Science Bldg., University of Arkansas, Fayetteville, AR 72701. Contributed equally (Soumen Nandy, Bhuvan Pathak).

## Abstract

Transient expression of CRISPR/Cas9 is an effective approach for limiting its activities and improving its precision in genome editing. Here, we describe the heat-shock inducible CRISPR/Cas9 system for controlled genome editing, and demonstrate its efficiency in the model crop, rice. Using a soybean heat-shock protein gene promoter and the rice *U3* promoter to express Cas9 and sgRNA, respectively, we developed the heat-shock (HS) inducible CRISPR/Cas9 system, and tested its efficacy in targeted mutagenesis. Two loci were targeted by transforming rice with HS-CRISPR/Cas9 vectors, and the presence of targeted mutations was determined before and after the HS treatment. We found only a low rate of targeted mutagenesis before HS, but an increased rate of mutagenesis after HS treatment among the transgenic lines. Specifically, only ∼11% of transformants showed characteristic insertions-deletions at the ambient room temperature, but a higher percentage (∼45%) of callus lines developed mutations after a few days of HS treatment. Analysis of regenerated plants harboring HS-CRISPR/Cas9 revealed that targeted mutagenesis was suppressed in the plants but induced by HS, which was detectable by Sanger sequencing after several weeks of HS treatments. Most importantly, the HS-induced mutations were transmitted to the progeny at a high rate, generating monoallelic and biallelic mutant lines that independently segregated from Cas9. Taken together, this work shows that HS-CRISPR/Cas9 is a controlled and reasonably efficient platform for genome editing, and therefore, a promising tool for limiting genome-wide off-target effects and improving the precision of genome editing.

**Significance Statement:** A method for the temporal control on gene editing based on the use of heat-shock induced expression of CRISPR/Cas9 is described, which was efficient in producing heritable mutations in the rice genome. We assume this method will be useful for targeting essential genes and improving the precision of CRISPR/Cas9.

## Introduction

The CRISPR/Cas9 system is an efficient tool for genome editing that is gaining popularity in both agricultural and medical biotechnology. It consists of two components: Cas9 nuclease and single-guide RNA (sgRNA) that form a complex (sgRNA:Cas9) and target sequences complementary to ∼20 nt spacer sequence in sgRNA, provided the NGG protospacer adjacent motif (PAM) is located at the *3’* end of the target sequence. Successful targeting by Cas9 results in a blunt double-stranded break (DSB), 3-nt upstream of the NGG motif (Cong et al., 2013; Jinek et al. 2012; Mali et al., 2013; Mojica et al. 2009), the repair of which by the cell leads to gene editing effects such as insertion-deletions (indels) and gene replacement (Jasin and Haber, 2016; Puchta et al. 1996; Rouet et al. 1994; Szostak et al. 1983; Waterworth et al. 2011). Similarly, CRISPR/Cas12a, an alternative gene editing tool, can be deployed on sequences ending with TTTN motifs (Endo et al., 2016; Schindele et al., 2018; Wang et al., 2017; Zetsche et al., 2015).

To improve gene editing efficiency, many different Cas9 expression systems have been described that mostly include developmental and constitutive gene promoters (Feng et al., 2018; Ma et al., 2016; Miki et al., 2018; Wang et al., 2015). In monocots, rice and maize ubiquitin promoters for Cas9 expression and *U3* or *U6* promoter for sgRNA expression are quite successful in creating targeted effects in the primary transformed (T0) plants (Lee et al., 2018; Wang et al., 2014; Xie and Yang, 2013). Previous studies have also shown that CRISPR/Cas9 effects could occur at a high rate during tissue culture or regeneration phases, leading to edited T0 lines that efficiently transmit the mutations to the next generation (Mikami et al., 2015; Zhang et al., 2014a; Srivastava et al., 2017). However, in these approaches, strong doses of sgRNA:Cas9 could persist far beyond the incidence of targeted gene editing, and provide the opportunity to mutagenize genome-wide off-target sites. Accordingly, off-targeting was found to be higher with higher doses of sgRNA:Cas9 in human cells, and ∼100x higher with constitutive-Cas9 as compared to transient-Cas9 in maize cells (Hsu et al., 2013; Pattanayak et al., 2013; Svitashev et al., 2015). The dose of sgRNA:Cas9 determines targeting efficiency; however, since mismatches between sgRNA spacer sequence and the target genomic sites are allowed at the PAM-distal end (Fu et al., 2013; Jinek et al., 2012; Lin et al., 2014; Liu et al., 2016), each sgRNA could potentially target numerous off-sites in the genome. Although, off-sites would generally be targeted at lower rates than the bona-fide target site, constitutive or tissue-specific expression systems would be more permissive to off-site mutations by providing strong doses of Cas9 for a longer than necessary period of time.

Off-target effects of CRISPR/Cas9 is a topic of intense investigation. Although, genetic segregation is an option for removing such mutations in many plant species, curbing off-target effects will be a better approach for developing high quality edited lines. Several approaches for improving the precision of gene editing have been described, *e.g*., high fidelity Cas9, split-Cas9, and ribonucleoprotein (RNP) Cas9 (Liang et al., 2017; Kleinstiver et al., 2016; Senturk et al., 2017; Svitashev et al., 2016; Wright et al., 2015). The use of RNPs has additional benefits in plant biotechnology as this DNA-free approach generates targeted mutations without incorporating foreign genes (Wolter and Puchta, 2017; Wolt et al., 2016). However, RNP approach in plants is faced with the difficulty of delivering the reagent in the cell wall bound compartments, and recovering the edited lines without selection in the tissue culture.

Here, we describe the use of inducible expression system for controlling CRISPR/Cas9 mutagenesis. Our rationale is to generate short phases of Cas9 expression in the tissue culture or regenerated plants for allowing targeted genome editing but keeping Cas9 suppressed at most other times until genetic segregation. In addition to helping reduce off-target effects, this temporal control on Cas9 could improve gene editing efficiencies by inducing Cas9 in phases conducive to gene editing, *e.g*., plant regeneration phase in the tissue culture (Zhang et al., 2014a; Srivastava et al., 2017), and enable conditional targeting to avoid lethal effects of mutations. Using heat-shock inducible promoter to express Cas9 and rice *U3* promoter for sgRNAs, we developed transformed lines of rice that essentially contained heat-shock (HS) controlled CRISPR/Cas9 system. By targeting genomic loci with paired sgRNAs, we determined the efficacy and efficiency of HS-CRISPR/Cas9 system in rice, the model cereal crop. Our analysis indicates that HS-CRISPR/Cas9 rarely induced mutations at the ambient room temperatures but efficiently created mutations upon heat-shock treatment in the callus and the regenerated plants. Notably, targeted mutations were transmitted to the progeny at a high rate and segregated independently from Cas9. In comparison with strong constitutive expression system consisting of rice Ubiquitin promoter (*RUBI*) to express Cas9 (Xie et al., 2015), HS-CRISPR/Cas9 created mutations at ∼50% rate. Sanger sequencing on predicted off-sites did not find mutations in either RUBI-or HS-CRISPR/Cas9 lines; however, to determine off-site targeting in the two systems, additional analysis with multiple sgRNAs and deep sequencing on whole genome and multiple off-sites will be needed. Overall, this study shows that HS-CRISPR/Cas9 is a controlled and efficient system for creating targeted mutagenesis, and therefore, a promising platform of improving gene editing that would be less permissive to off-target effects.

## Results

### Heat-shock induced CRISPR/Cas9 mutagenesis in the rice callus

We used soybean heat-shock protein 17.5E (*HSP17.5E*) gene promoter to express the humanized *Streptococcus pyogenes* Cas9 (SpCas9), and the tRNA-processing system to express two sgRNAs by the rice snoRNA *U3* promoter (Czarnecka et al., 1992; Xie et al., 2015; Fig. 1a, b). The motivation to use *HSP17.5E* promoter was based on its efficacy in controlling Cre-*lox* recombination in the tissue-culture derived rice plants and seedlings. Previously, we showed that a simple heat treatment of 42°C for 3 h led to efficient Cre-*lox* mediated excision of marker gene in rice seedlings and the inheritance of the marker-free locus by their progeny (Nandy and Srivastava, 2012). We chose previously tested target loci and sgRNAs for this study that include rice *Phytoene Desaturase* gene (*OsPDS*) and the *β–Glucuronidase* transgene inserted in rice genome (Srivastava et al., 2017). For *GUS* targeting, a well-characterized transgenic line, B1 (cv. Nipponbare), that harbors a single-copy of the *GUS* gene driven by maize ubiquitin promoter (*Ubi*), and for *PDS* targeting, non-transgenic Nipponbare were transformed. The resulting hygromycin-resistant calli were maintained and regenerated at the ambient room temperature. Randomly sampled calli cultures were transferred to fresh media plate for heat-shock (HS) treatment and the screening of mutations before HS (pre-HS) and after HS (post-HS) at the two targeted sites in each gene. One of the 12 PDS calli was found to contain pre-HS mutations at the predicted DSB sites (Fig. 1c-d; Table 1), indicating targeting due to high background CRISPR/Cas9 activity in this line. Similarly, one of the 6 GUS calli was found to contain pre-HS mutations at sg1 target (Fig. 2a; Table 1). Next, the callus samples were subjected to heat-shock (HS) treatment for 3 hours and returned to ambient room temperature for further growth. After 5 – 7 days, freshly grown tissue from each callus culture was analyzed. However, since calli could contain multiple independent mutations, Sanger sequencing would generate overlapping peaks downstream of the predicted DSB sites. Accordingly, overlapping traces in the sequencing spectra were found in 5 of the *PDS* lines and 3 *GUS* lines, indicating mosaic pattern of mutations due to HS-CRISPR/Cas9 activity (Fig. 1c-d, 2b; Table 1). To verify these mutations, TA cloning and colony sequencing was done on a subset of samples representing *PDS* sg1 and sg2 targets, and *GUS* sg2 targets. Characteristic indels at the predicted DSB sites were found in >1 clone in each sample, confirming CRISPR/Cas9 mediated mutagenesis (Fig. 1e-f; 2d). In conclusion, although, occurrence of pre-HS mutations in rice calli cannot be ruled out, *HSP17.5E-Cas9* is effective in creating HS-induced targeted mutations. With the paired sgRNAs targeting a gene, HS-CRISPR/Cas9 generated HS-induced mutations in ∼40 – 50% of the transformants (**Table S1**). All callus cultures were subjected to plant regeneration; however, PDS cultures mostly appeared non-embryogenic, while GUS cultures regenerated plants. Therefore, all subsequent work was done with HS-CRISPR/Cas9 targeting the *GUS* transgene.

**Figure 1:**
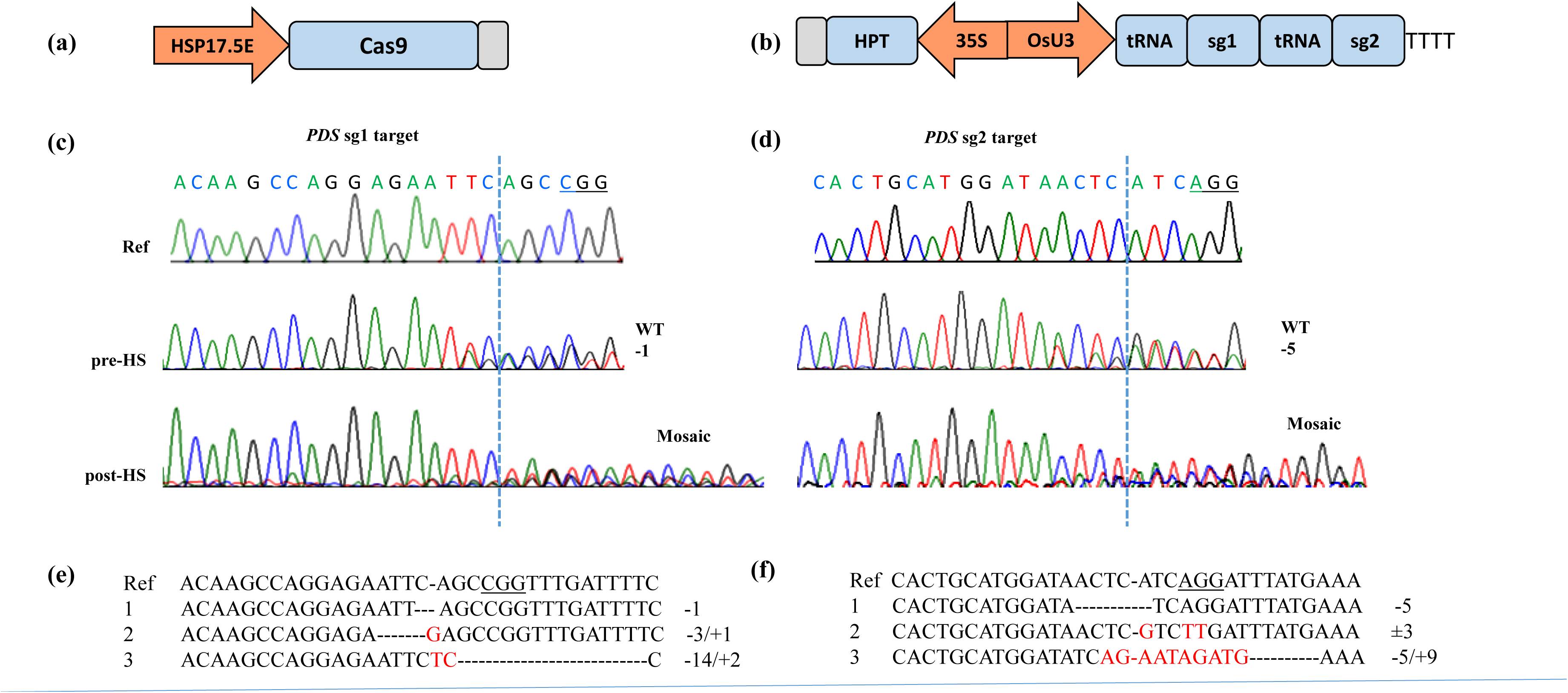
Efficacy of heat-shock (HS) inducible CRISPR/Cas9 on rice *Phytoene Desaturase* (*PDS*) gene. **(a)** HS-Cas9 expression construct consisting of soybean heat-shock protein 17.5E (HSP17.5E) gene promoter and the *Streptococcus pyogenes* Cas9 coding sequence; **(b)** standard sgRNA construct consisting of rice sno U3 promoter expressing a pair of sgRNAs via tRNA processing mechanism. For plant selection, hygromycin resistance gene consisting of 35S promoter and hygromycin phosphotransferase (HPT) gene was included in the construct. Pol III terminator is shown as TTT, and gray bars represent *nos 3’* terminators; (**c-d**) Sequencing spectra of *OsPDS* target sites (PAM underlined) on non-transgenic (ref.) and representative HS-CRISPR/Cas9-transformed callus lines, before heat-shock (pre-HS) or after a few days of HS treatment (post-HS). Targeted mutations are indicated by two (heterozygous) or multiple overlapping sequence traces (mosaic) near the predicted double-stranded break (DSB) site (dotted line) in the spectra; **(e-f)** Alignments of the reference sequence with mutant reads as identified by CRISP-ID tool or TA cloning. Insertion-deletions (indels) are indicated by red fonts and dashed lines. Number of insertions or deletions is also indicated. PAM site (underlined) and predicted DSB sites (-) are indicated in the reference sequences.

**Figure 2:**
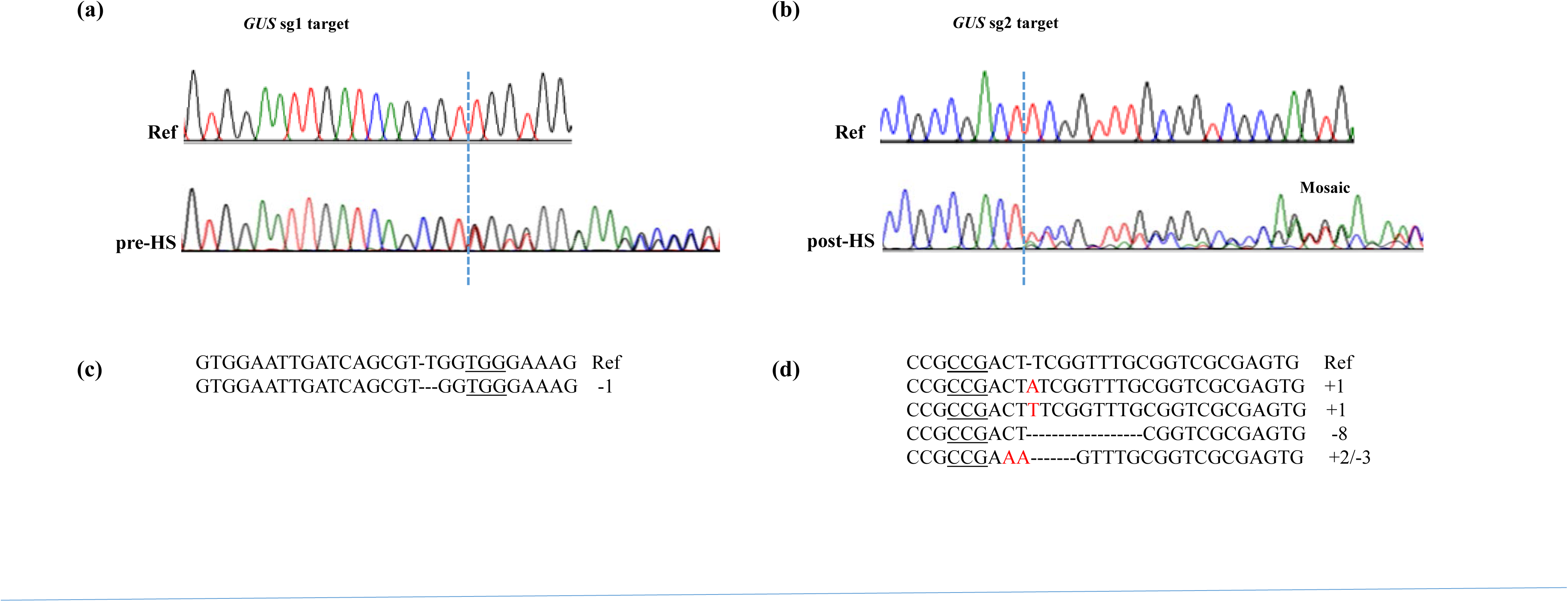
Efficacy of HS-CRISPR/Cas9 on the *GUS* transgene located in rice genome. (**a-b**) Sequencing spectra of the *GUS* target sequences from the untargeted *GUS* line (ref.) and the targeted callus lines, before heat-shock (pre-HS) or subsequent to HS treatment (post-HS). Dotted vertical lines represent the predicted DSB sites. Overlapping sequence traces in the spectra indicate mutations; (**c-d**) Mutations in the spectra identified as by CRISP-ID tool (c) or TA cloning (d). Dashes indicate deletions, and red letters indicate insertions. Number of insertions-deletions in each sequence is indicated. PAM site (underlined) and predicted DSB sites (-) are indicated in the reference sequences.

**Table 1:** HS-CRISPR/Cas9 activity in rice callus.

| Exp. | Target | No. of calli analyzed | Pre-HS mutations<sup>1</sup> |  | Post-HS mutations<sup>2</sup> |  | Efficiency<sup>3</sup> |  |
| --- | --- | --- | --- | --- | --- | --- | --- | --- |
|  |  |  | Sg1 | Sg2 | Sg1 | Sg2 | Sg1 | Sg2 |
| 1 | *OsPDS* | 12 | 1 | 1 | 3 | 5 | 25 | 41.6 |
| 2 | *GUS* | 6 | - | 1 | 1 | 3 | 16.6 | 50 |

<sup>1</sup>No. of callus lines showing mutations without heat-shock treatments
<sup>2</sup>No. of callus lines showing mutations subsequent to heat-shock treatments
<sup>3</sup>Percent transformants showing mutations after heat-shock treatments

### Heat-shock induced targeting in T0 plants

Twenty regenerated plants (T0) expressing HS-CRISPR/Cas9 against *GUS* gene were obtained from two experiments. One – three leaf samples from each plant were subjected to PCR and Sanger sequencing at the targeted sites. Two of the T0 plants (#9 and #12) were found to harbor biallelic homozygous or heterozygous mutations at the sg2 target, indicating misregulation of Cas9 in these plants (Fig. 3). The rest did not show mutations at either site (Table 2). Subsequently, T0 plants were given two rounds of HS treatment by transferring them to 42°C incubator for 3 hours and repeating the treatment after ∼20 h of rest at the room temperature, and subsequently transplanted in the soil and grown in the greenhouse. After ∼4 wks of HS treatment, at the young vegetative stage, target site analysis by PCR and sequencing was conducted in 2 – 3 leaf samples harvested from different tillers. No detectable targeting was found in any of the samples except those derived from T0#9 and #12; although, a baseline secondary sequence was detected in the sequencing spectra of a few lines, indicating a low rate of HS-induced mutations (Fig. 4, S1; Table 2). This observation corroborated with histochemical GUS staining as these plants progressively lost GUS activity. For example, T0#1 showed strong GUS staining in the leaf cuttings taken from the young plant but no staining in the leaves collected from the flowering plant, while T0#2 continued to show strong GUS staining and lacked detectable mutations (Table 2). As expected, T0 #9 and #12 that harbored biallelic mutations did not display GUS staining in the leaves derived from the vegetative or flowering stages of the plant. These observations are analogous to our work with HS Cre-*lox* system, in which, rice seedlings harboring *HS-Cre* showed progressive recombination in heat-shocked plants, and transmitted the recombined locus to the next generation (Nandy and Srivastava, 2012). Taken together, HS-induced gene editing effects likely occurred in the early cell lineages and established in the plant through cell division.

**Figure 3:**
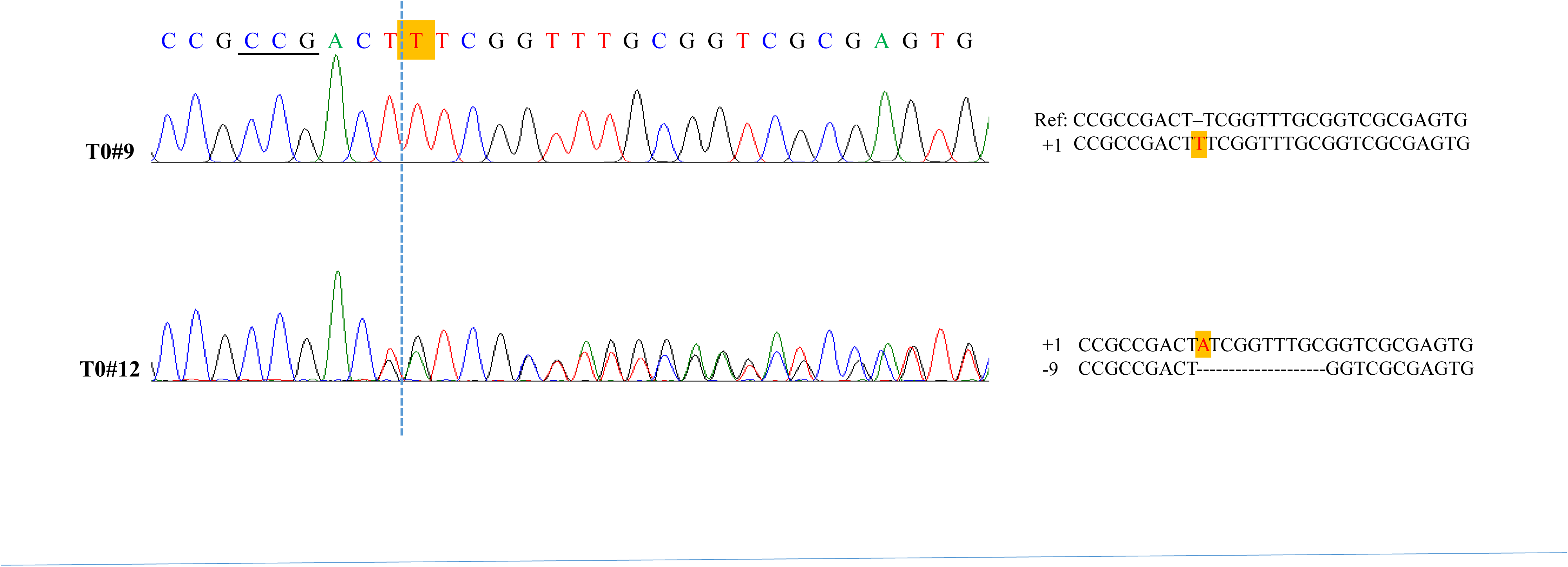
Sequencing of *GUS* sg2 target site in T0 plants #9 and #12 harboring HS-CRISPR/Cas9. Mutation types are shown adjacent to each spectra along with the reference sequence. Dashed vertical line indicates the predicted DSB site. PAM site is underlined. Shaded red letter indicate insertions, and dashes indicate deletions. The two sequences in T0#12 were separated using the CRISP ID tool.

**Figure 4:**
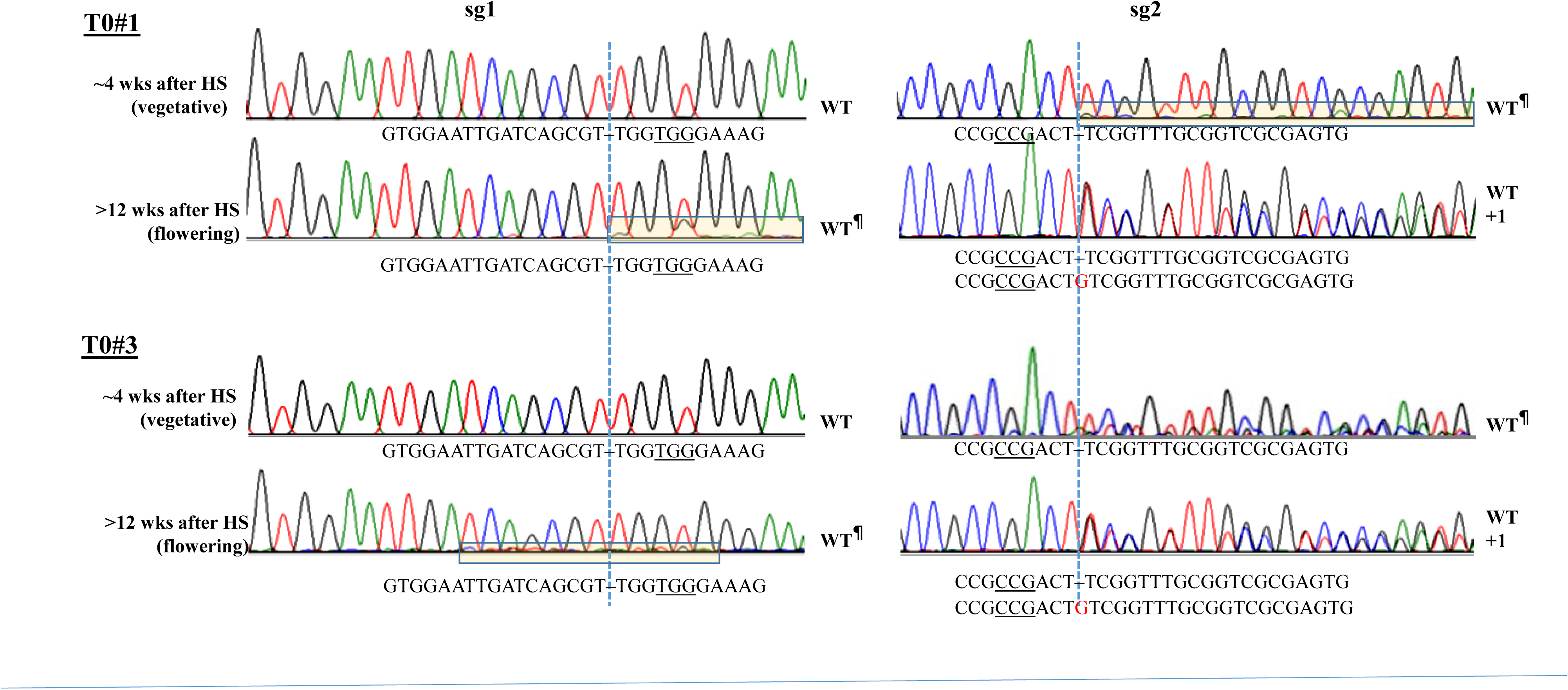
Genotyping of T0 plants #1 and #3 at *GUS* sg1 and sg2 sites by PCR-sequencing at two growth stages, ∼4 weeks after heat-shock (HS) or vegetative stage and ∼12 weeks after HS or flowering stage. Mutation types are shown below each sequencing spectra with PAM sequence underlined. The predicted DSB sites are indicated by the vertical line. The baseline secondary sequence traces in the spectra are boxed, indicating a low rate of mutations in largely wild-type samples (WT^¶^). The spectra containing 2 overlapping sequences were analyzed by CRISP-ID tool to identify monoallelic +1 mutations in the two plants.

**Table 2:** Characterization of T0 Plants transformed with HS-CRISPR/Cas9 targeting GUS gene.

| Line | GUS staining<sup>#</sup> |  | Cas9 expression |  | Sg1 | Sg2 |
| --- | --- | --- | --- | --- | --- | --- |
|  | Y | O | Fold-induction by HS | % RUBI-Cas9 |  |  |
| 1 | + | - | 5.7 | 0.03 | WT | WT<sup>¶</sup> |
| 2 | + | + | 0.35<sup>†</sup> | 0.07 | WT | WT |
| 3 | ± | - | 2.5 | 0.13 | WT | WT<sup>¶</sup> |
| 4 | + | + | 7.3 | 0.02 | WT | WT |
| 5 | + | + | 94 | 0.03 | WT | WT |
| 6 | + | + | - | - | WT | WT |
| 7 | + | ± | - | - | WT<sup>¶</sup> | WT<sup>¶</sup> |
| 8 | + | + | - | - | WT | WT |
| 9 | - | - | - | - | WT | Biallelic |
| 10 | + | + | 0.45<sup>†</sup> | 0.2 | WT | WT |
| 11 | + | + | - | - | WT | WT |
| 12 | - | - | 53x | 5.96 | WT | Biallelic |
| 13 | + | ± | - |  | WT | WT<sup>¶</sup> |
| 14 | + | + | 1<sup>‡</sup> | 16.96 | WT | WT |
| 15 | + | + | 2.2 | - | WT | WT |
| 16 | + | + | - | - | WT | WT |
| 17 | + | + | - | - | WT | WT |
| 18 | + | - | 6.4 | 0.09 | WT | WT |
| 19 | + | ± | 6.2 | 0.02 | WT<sup>¶</sup> | WT |
| 20 | + | + | 4.1 | 0.03 | WT | WT |
| RUBI-1 | - | - | - | 100 | Biallelic | Biallelic |
| RUBI-2 | + | + | - | 100 | - | - |
| RUBI-3 | + | + | - | 50 | - | - |

<sup>#</sup>Histochemical staining of leaf cuttings from young vegetative (Y) or older flowering (O) plants.
<sup>†</sup>Silenced Cas9 lines
<sup>‡</sup>Overexpression Cas9 lines
<sup>¶</sup>Baseline secondary sequence trace in the sequencing spectra (see Fig. S1).

T0 plants #1, #2, #3 were the first to flower and set seeds. These plants were analyzed again for the presence of mutations at the target sites (>12 wks after HS treatment). As shown in Fig. 4, these plants at the flowering stage showed minor targeting at sg1 site; whereas, a clear monoallelic targeting was observed at sg2 site. Since, low rate of mutagenesis at sg2 was detected in these plants at the young vegetative stage (baseline secondary sequence in the spectra) (Fig. 4), these monoallelic mutations were most likely induced early in the plant. T0#2, however, did not show mutations in any analyzed tissue, and later was found to contain non-inducible, possibly silenced Cas9 (described below).

Cas9 gene expression was analyzed in a subset of T0 plants and compared with non-transgenic wild-type and constitutive Cas9 lines using real-time quantitative PCR. Of 12 plants, 9 showed increase in Cas9 expression (2 – 94x) upon HS over their respective room temperature (RT) values (Fig. 5a; Table 2). Two T0 plants (#2, #10) appeared to be silenced as the relative Cas9 expression in these plants did not increase by HS treatment, whereas one (#14) showed equally high expression at RT and HS, which was 16x higher than the constitutive-Cas9 lines (Table 2). Three constitutive-Cas9 lines expressing RUBI-CRISPR/Cas9 (RUBI-1, 2, 3) were included in the analysis, each of which showed strong relative expression, and one of them (RUBI-1) harbored targeted mutations in the *GUS* gene (Table 2). In comparison to RUBI-Cas9 lines, Cas9 expression was three orders of magnitude lower in HS-Cas9 lines, which could be induced ∼100-fold by HS (Fig. 5b; Table 2).

**Figure 5:**
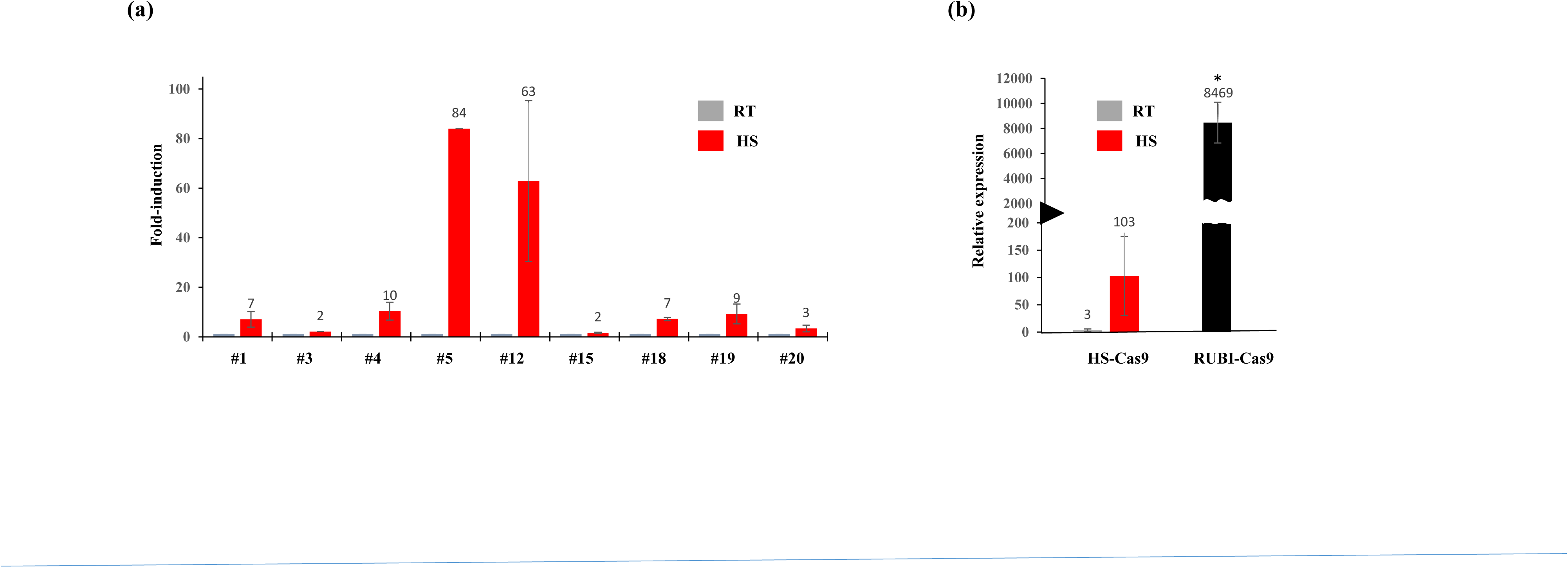
Cas9 expression analysis. **(a)** Fold-induction of Cas9 in T0 plants by heat-shock (HS) treatment (3 h exposure to 42°C) as compared to background room-temperature (RT) values; **(b)** Relative expression of Cas9 in HS-Cas9 lines and the constitutive RUBI-Cas9 lines. The expression in HS-Cas9 lines was calculated at RT and upon HS. The average of 8 HS-Cas9 lines and 3 RUBI-Cas9 lines is shown with standard errors (*p-value <0.001).

### Inheritance of targeted mutations by the progeny

T0#1, #2, and #3 were the first to flower and produce seeds. However, Cas9 was silent in #2; accordingly, targeting was undetectable in this plant (Table 2). Therefore, T0#1 and #3 were selected for the progeny analysis. As shown in Table 2, these plants, at the young vegetative stages, showed relatively high GUS activity compared to that in the flowering stages, presumably due to the division of cells harboring mutations in the *GUS* gene. Sequencing of sg1 and sg2 sites in these plants at the vegetative and flowering stages detected targeted mutagenesis by CRISPR/Cas9 in one or both sites (Fig. 4).

Twenty-four seeds derived from T0#1 parent and 30 seeds from T0#3 parent were germinated. When their coleoptiles were fully emerged, these seedlings were subjected to 2 – 3 rounds of HS treatment. Therefore, *de novo* targeting could occur in Cas9+ lines. Histochemical GUS staining of these seedlings (∼2 wks after germination) showed either presence (+), absence (-) or mosaic pattern (±) of staining. As expected, *Cas9* independently segregated in the population, and a few null-segregants were identified in the two populations (Table 3). A subset of 16 T1 plants derived from T0#1 were subjected to PCR/sequencing at sg1 and/or sg2 sites. At sg1 site, eleven contained monoallelic (68.7%) and two biallelic mutations (6.2%), while at sg2 site, nine contained monoallelic (56.2%) and two biallelic (6.2%) mutations (Table 3**; Table S2**). Analysis of 25 T0#3 progeny, on the other hand, revealed monoallelic and biallelic mutations at sg1 site in eighteen (72%) and two (8%), respectively, while at sg2 only monoallelic mutants (96%) were found (Table 3**; Table S3**). The remaining inherited WT alleles. The analysis of mutant reads revealed 4 – 5 types of mutations among T0#1 progeny but only one type at each site among T0#3 progeny (Fig. 6). The abundance of one type of mutation in each population indicates high rate of inheritance, which was confirmed by three Cas9 null-segregants in each population that harbored mutations at sg1 and/or sg2 sites (Fig. 6c-d**; Table S2, S3**). The detection of only one type of mutation among #3 progeny raises the question whether this line is derived from *HS-Cas9* activity induced by the tissue culture. However, since the analysis of 3 different leaf samples of T0#3 plant detected only WT sg1 site (Fig. 4), the observed mutations are likely established in germline after HS treatment in this plant.

**Figure 6:**
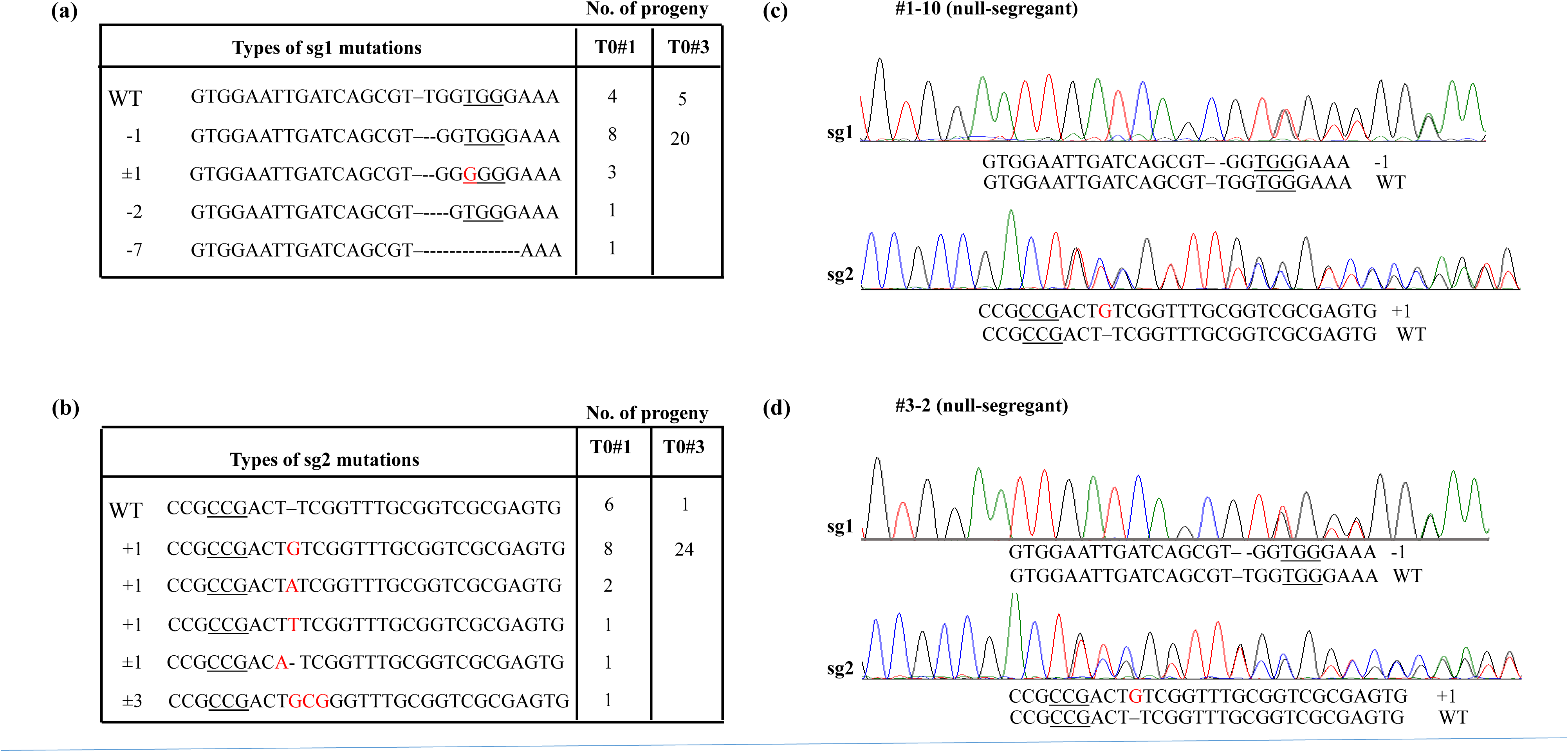
Inheritance of HS-CRISPR/Cas9 induced mutations by the progeny of T0#1 and #3. (**a-b**) Number of T1 plants harboring monoallelic ot biallelic indels at *GUS* sg1 and sg2 target sites. Indels in each mutations are shown as dashes and red letters; **(c-d)** Inheritance of mutations in the two Cas9 null-segregants harboring monoallelic mutations at sg1 and sg2 sites. The sequence reads as identified by separating overlapping reads by CRISP-ID tool and their alignments are shown below each spectra. Insertion and deletion are shown by red letter or dashes. PAM is underlined.

**Table 3:**
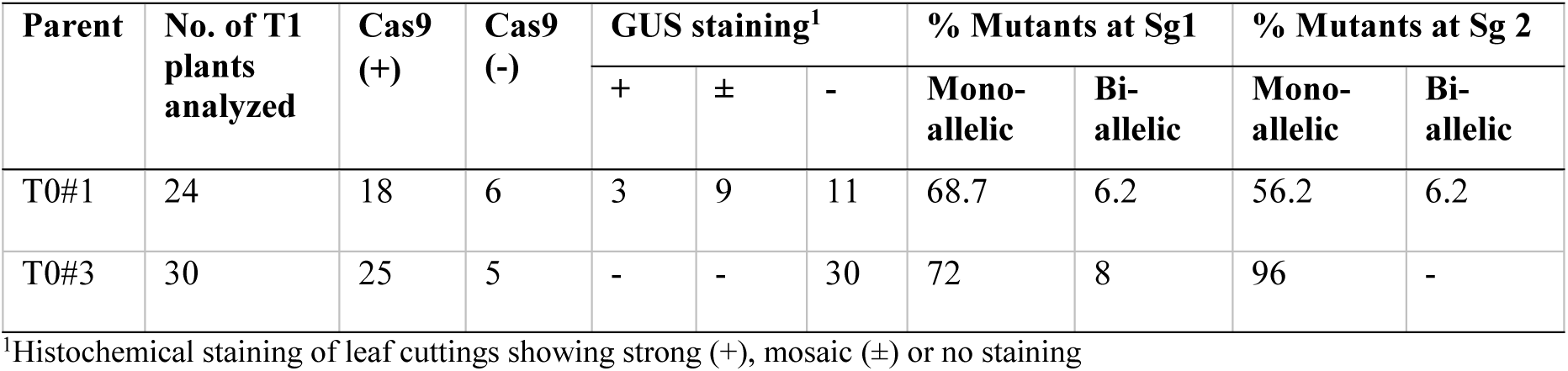
Analysis of T1 progeny.

| Parent | No. of T1 plants analyzed | Cas9 (+) | Cas9 (-) | GUS staining<sup>1</sup> |  |  | % Mutants at Sg1 |  | % Mutants at Sg 2 |  |
| --- | --- | --- | --- | --- | --- | --- | --- | --- | --- | --- |
|  |  |  |  | + | ± | - | Mono-allelic | Bi-allelic | Mono-allelic | Bi-allelic |
| T0#1 | 24 | 18 | 6 | 3 | 9 | 11 | 68.7 | 6.2 | 56.2 | 6.2 |
| T0#3 | 30 | 25 | 5 | - | - | 30 | 72 | 8 | 96 | - |

<sup>1</sup>Histochemical staining of leaf cuttings showing strong (+), mosaic (±) or no staining

## Discussion

The CRISPR/Cas9 system shows high efficiency targeting in plants and animals, and is often described as a precise system that generates limited or undetectable off-target effects in plants (Feng et al., 2018; Lee et al., 2018; Tang et al., 2018). However, since the mechanism of targeting is based on a short-stretch of sequence complementarity and presence of a trinucleotide PAM (NGG) (Jinek et al., 2012), and since mismatches are tolerated at the PAM-distal end, numerous sites in a complex genome could potentially fall within the scope of CRISPR/Cas9 targeting. Further, sequences ending with non-canonical PAMs such as NAG can also be targeted by Cas9 (Zhang et al., 2014b), and while chromatin structure plays a marginal role in targeting, secondary structures in the target DNA and sgRNA could allow significant pairing, in spite of mismatches at the PAM end (Lin et al., 2014). Therefore, current bioinformatics tools are limited in their ability in accurately predicting off-target sites of designed sgRNAs. In the mammalian cells, high concentrations of sgRNA:Cas9 reportedly induced high rate of off-target mutations (Hsu et al., 2013; Pattanayak et al., 2013). Similarly, in plants, off-targeting increased 100-fold with constitutive-Cas9, in comparison to transient Cas9 (Svitashev et al., 2015).

In plants, ribonucleoprotein Cas9 (RNP) has been used as an effective transient expression system (Liang et al., 2017; Svitashev et al., 2016). However, the efficiency of RNP in plant cells is impacted by the difficulty in delivering it into the cell wall bounded compartments and isolating the edited lines in the selection-free transformation system (Yin et al., 2017). Inducible expression systems can be argued as more versatile transient expression systems, provided they generate low or undetectable background expression and high induced expression. Heat-shock promoters meet these criteria as they have been successfully used in applications where their proper regulation was critical, *e.g.,* controlling Cre-*lox* recombination or nuclease activity for marker excision (Khattri et al., 2011; Lloyd et al., 2005; Nandy et al., 2015; Nandy and Srivastava, 2012; Zhang et al., 2003).

Here, we describe the use of heat-shock (HS)-CRISPR/Cas9 system consisting of HS inducible expression of Cas9 and the standard *U3* promoter for sgRNA expression. We found that HS-CRISPR/Cas9 at the room temperature was suppressed in rice tissue culture and regenerated plants as mutations in the targeted sites were undetectable in most lines analyzed in this study. However, upon HS treatment, characteristic CRISPR/Cas9 mutations at the two *OsPDS* target sites were found in 25 – 41% of calli, and at the two *GUS* target sites in 16 – 50% of the calli (Table 1). It is well known that targeting efficiency varies between genomic sites. In our work with constitutive (RUBI)-CRISPR/Cas9, the two *OsPDS* sites were targeted among ∼75% of transformants and the *GUS* sg2 site among 42% (**Table S4**). Therefore, relative targeting efficiency of HS-Cas9 with one or two rounds of HS treatments appears to be ∼50% of the constitutive-Cas9. Whether this efficiency could be further improved by additional HS treatments is yet to be determined. However, the two Cas9 expression systems could not be compared in T0 plants as HS-induced mutations in the plants are evident only as chimeric rare mutations, indicated by the baseline secondary sequence in the sequence spectra (**Fig. S1**). However, in plants, inheritance rate is the most important criteria of gene editing efficiency. We show that HS-induced mutations in T0 plants were transmitted to the progeny at a high rate and segregated independently from *Cas9* (Table 3). Further, our data reflects on the efficiency of HS-CRISPR/Cas9 is inducing mutations in the meristem, leading to the mutant cell lineage in the somatic tissue and the germline, which explains the high frequency of one type of mutation observed in the progeny, especially, in the T1 population of T0#3 (Fig. 6).

We also tested 3 potential off-target sites of *GUS* sgRNA1, identified by Cas-OFFinder or GGGenome tool (Bae et al., 2014; www.gggenome.dbcls.jp; Table S5) in 15 independent edited plants (T0 and T1) by PCR and Sanger sequencing, and included wild-type Nipponbare, Parent B1, and RUBI-CRISPR/Cas9 lines for comparison. No indels were detected in any of the lines at the three off-target sites; although, in one of the sites (#2, Table S5) that is located in the intergenic region on Chromosomes 1, single-nucleotide polymorphisms (SNPs) were detected in 4 T1 plants (**Fig. S2**). However, since indels are more consistent with the CRISPR/Cas9 mutagenesis (Tang et al., 2018), the observed SNPs are more likely the effects of tissue culture on the rice genome. To clearly compare and contrast the off-targeting frequencies of HS-and RUBI-CRISPR/Cas9 systems, deeper analysis with multiple sgRNAs will be necessary. This study, however, focused on the efficiency and efficacy of HS-CRISPR/Cas9, and demonstrated the utility of inducible expression systems in plant genome editing.

Several drug-inducible gene editing systems have been described for human cells (Dow et al., 2015; Nihogaki et al., 2018), but heat-inducible Cas9 has so far been used only in *Caenorhabditis elegans* (Li et al., 2015; Liu et al., 2014). In addition to their potential in curbing off-target effects, inducible expression systems could confer spatio-temporal control on gene editing, which can simplify editing of essential genes, avoid lethality by activating Cas9 at specific developmental stage, and improve gene editing efficiency by inducing Cas9 in the repair-competent cells. Use of heat-inducible expression system could also leverage improved Cas9 activity by heat-shock, leading to higher rates of mutagenesis (LeBlanc et al., 2018). Additionally, heat-shock was found to enhance sgRNA levels (**Fig. S3**), which could improve gene editing efficiency, if sgRNA is limiting. Although, the molecular basis of heat-induction of sgRNAs is not clear, a similar observation was made in Arabidopsis by LeBlanc et al. (2018).

In summary, we demonstrate HS-inducible CRISPR/Cas9 system is regulated properly in rice by keeping Cas9 suppressed at the ambient room temperature, and activating Cas9 by heat-shock treatment. The heat-shock induced genome editing is reasonably efficient at producing heritable targeted mutations as demonstrated by targeting two loci in rice. Therefore, HS-inducible CRISPR/Cas9 could prove to be an important genome editing tool in plant biotechnology that can provide temporal control towards improving the precision of CRISPR/Cas9 activities. This expression platform could also be used for the temporal control of other gene editing tools such as CRISPR/Cas12a

## Acknowledgements

The vectors pRGE32 and pGTR that were used for construction of HS-Cas9 and sgRNA constructs were donated by Yinong Yang and obtained from Addgene.com. Technical support in making paired sgRNA constructs was provided by Jamie Underwood, and greenhouse support was provided by Jessica Kivett. This project is supported by the ABI-Arkansas Division of Agriculture and the USDA-NIFA grant 2017-38821-26412.

## Experimental Procedures

### DNA Constructs and Plant Transformation

The *Cas9* coding sequence was PCR amplified from pRGE32 (Addgene #63159) using primers (Table S4) laced with specific restriction enzyme sites and cloned between soybean *HSP17.5E* gene promoter (GenBank accession no. M28070) and nopaline synthase terminator (*nos 3’*) in a pUC19 vector backbone. The sgRNA vectors were made in pRGE32 backbone using the protocol of Xie et al. (2015) and the sgRNA spacer sequences were selected using CRISPR RGEN tool (http://www.rgenome.net/cas-designer; Park et al. 2015). The resulting *GUS* (GenBank accession no. AF485783) and *OsPDS* (Os03g08570) sgRNA constructs were PCR amplified with primers shown in Table S5 and cloned into a vector harboring 35S promoter driven hygromycin phosphotransferase (*HPT*) gene. All vectors were verified by sequencing. B1 transgenic line (*cv.* Nipponbare), which has been described by Nandy and Srivastava (2012) or wild-type Nipponabare was used for transformation. B1 contains a single-copy of *GUS* gene controlled by maize ubiquitin-1 gene promoter. The GUS activity was verified by staining endosperms using the GUS staining solution described by Jefferson (1987). The embryogenic callus obtained from mature seeds of the homozygous B1 line was used for all transformations. All transformations were done by the gene gun (PDS1000, Bio-Rad Inc.) based DNA delivery of the Cas9 and the sgRNA vectors (Fig. 1a). The transformed calli were isolated on hygromycin (50 mg/l) containing media. All tissue culture and regeneration in this study was done using the method of Nishimura et al. (2006).

### Heat-shock Treatments

Freshly plated calli, rooted regenerated plants in glass tubes or ∼1 wk old seedlings on MS/2 plates were subjected to heat-shock (HS) treatment by transferring them to pre-heated 42°C incubator. The Petri dishes containing calli or germinating seedlings and glass tubes containing regenerated plants were laid on their sides between pre-heated metal plates. After 3 hours, plates or tubes were returned to tissue culture chamber set at 25°C for further growth. Tissues were harvested after a few days for genotyping by PCR and sequencing.

### DNA Extraction, PCR and Sequencing

Genomic DNA isolated from callus, regenerated plants or seedlings was used for polymerase chain reaction (PCR) using primers spanning the target sites (Table S5) or predicted off-target sites (Table S6). The PCR products were resolved on agarose gel and extracted using GeneJET Gel Extraction Kit (Thermo Scientific, USA) for sequencing from both ends using forward and reverse primers by the Sanger Sequencing method at Eurofins Genomics USA (www.eurofinsgenomics.com). Selected PCR amplicons were cloned into pCR2.1 vector using TA cloning kit (Thermo-Fisher Scientific, NY) as per manufacturer’s instructions. Randomly picked fifteen to twenty colonies were verified for the insert by PCR using amplicon-specific primers and sequenced at Eurofins Genomics USA. The sequence traces (ABI files) were analyzed on Sequence Scanner 2 software (Applied Biosystems Inc.) and aligned with the reference sequences using CLUSTAL-Omega multiple sequence alignment tool. The overlapping sequence traces arising from heterozygous alleles or chimeric samples were separated using the CRISP-ID tool (Dehairs et al. 2016).

### Gene Expression Analysis

Young developing leaves were collected from the same tiller and incubated at room temperature (25°C) or 42°C for 3 hours for the control and heat-shock treatments, respectively. The total RNA was isolated from 100 mg samples using QIAGEN RNeasy plant mini kit (Qiagen, Valencia, CA), and treated with RNase-Free RQ1 DNase (Promega, San Luis Obispo, CA), and quantified using NanoDrop 2000 (Thermo Fisher Scientific, NY). The expression analysis on Cas9 and sgRNAs was performed on 25 ng of RNA using Superscript III Platinum SYBR green one step qRT-PCR (Life Technologies, Grand Island, NY) in the CFX96 Real Time PCR Detection system (Bio-Rad, Hercules, CA). The values were normalized against the rice ubiquitin gene, and the relative expression to the non-transgenic control was calculated using 2 ^ΔΔCt^ (Livak and Schmittgen, 2001) method. Standard errors of three to six biological replicates were calculated. Each biological replicate was repeated two times for the analysis. Student *t* test (unpaired) was used to determine the *p*-value. Primers used in qRT-PCR are given in Table S6.

---

**Figure S1:** Representative sequence spectra with baseline secondary sequence trace (boxed area) indicating a low rate of mutagenesis induced by HS-CRISPR/Cas9 activity. The target sites with PAM (underlined) are shown above each spectra.

**Figure S2:** Sequence alignments of predicted *GUS* off-target site obtained by PCR-sequencing from GUS line B1 and HS-CRISPR/Cas9 T1 plants. S1 and S2 refer to overlapping sequence traces in the ABI files. Single-nucleotide polymorphisms (SNP) are shown as red letters. sgRNA spacer sequence and the reference Nipponbare sequences are shown. Small blue letters in the reference sequences indicate mismatches with sgRNA spacer.

**Figure S3:** sgRNA expression analysis by real-time quantitative PCR in HS-CRISPR/Cas9 lines. Relative expression at room temperature (RT) and upon heat-shock (HS) at 42°C for 3 h. Average of 6 independent HS-CRISPR/Cas9 lines is shown as log10 transformed values relative to WT. Statistical differences (a, b) were determined by Student *t* test.

